# BEAST 2.5: An Advanced Software Platform for Bayesian Evolutionary Analysis

**DOI:** 10.1101/474296

**Authors:** Remco Bouckaert, Timothy G. Vaughan, Joëlle Barido-Sottani, Sebastián Duchêne, Mathieu Fourment, Alexandra Gavryushkina, Joseph Heled, Graham Jones, Denise Kühnert, Nicola De Maio, Michael Matschiner, Fábio K. Mendes, Nicola F. Müller, Huw Ogilvie, Louis du Plessis, Alex Popinga, Andrew Rambaut, David Rasmussen, Igor Siveroni, Marc A. Suchard, Chieh-Hsi Wu, Dong Xie, Chi Zhang, Tanja Stadler, Alexei J. Drummond

## Abstract

Elaboration of Bayesian phylogenetic inference methods has continued at pace in recent years with major new advances in nearly all aspects of the joint modelling of evolutionary data. It is increasingly appreciated that some evolutionary questions can only be adequately answered by combining evidence from multiple independent sources of data, including genome sequences, sampling dates, phenotypic data, radiocarbon dates, fossil occurrences, and biogeographic range information among others. Including all relevant data into a single joint model is very challenging both conceptually and computationally. Advanced computational software packages that allow robust development of compatible (sub-)models which can be composed into a full model hierarchy have played a key role in these developments.

Developing such software frameworks is increasingly a major scientific activity in its own right, and comes with specific challenges, from practical software design, development and engineering challenges to statistical and conceptual modelling challenges. BEAST 2 is one such computational software platform, and was first announced over 4 years ago. Here we describe a series of major new developments in the BEAST 2 core platform and model hierarchy that have occurred since the first release of the software, culminating in the recent 2.5 release.

**Author summary:** Bayesian phylogenetic inference methods have undergone considerable development in recent years, and joint modelling of rich evolutionary data, including genomes, phenotypes and fossil occurrences is increasingly common. Advanced computational software packages that allow robust development of compatible (sub-)models which can be composed into a full model hierarchy have played a key role in these developments. Developing scientific software is increasingly crucial to advancement in many fields of biology. The challenges range from practical software development and engineering, distributed team coordination, conceptual development and statistical modelling, to validation and testing. BEAST 2 is one such computational software platform for phylogenetics, population genetics and phylodynamics, and was first announced over 4 years ago. Here we describe the full range of new tools and models available on the BEAST 2.5 platform, which expand joint evolutionary inference in many new directions, especially for joint inference over multiple data types, non-tree models and complex phylodynamics.

## Introduction

Bayesian Evolutionary Analysis by Sampling Trees (BEAST) is a software package for performing Bayesian phylogenetic and phylodynamic analyses. BEAST samples from the posterior distribution of trees (or networks) and parameters given the input data using the Markov chain Monte Carlo (MCMC) algorihtm. Four years ago, BEAST 2 [1, 2] was published as a complete rewrite of the original BEAST software. A main goal of this rewrite was to develop a more modular software framework, one that could be easily extended by third parties. The software platform is comprised of various standalone programs including BEAUti (a graphical user interface [GUI] for setting up an analysis), BEAST to run MCMC analysis, and post processing tools such as LogAnalyser, LogCombiner, TreeAnnotator, DensiTree [3], as well as a package manager.

Shortly after its release, a number of packages were added, such as MASTER for simulating stochastic population dynamics models [4], MultiTypeTree for inferring structured coalescent models [5], RBS for reversible jump across substitution models [6], SNAPP for multi species coalescent over SNP data [7], subst-bma for Bayesian model averaging over site models [8], and BDSKY for the birth-death skyline tree model [9]. All these packages have been very popular on their own right, and since the initial release of BEAST 2 a large amount of functionality and packages have been added, showing the success of the approach. In this paper, we summarize the significant advances that have been made.

## What is BEAST?

BEAST is a package for conducting Bayesian phylogenetic inference using MCMC. At its core are rooted time trees (or time networks in latest developments), which can be inferred from multiple sources of data. BEAST supports sequence data for nucleotides, amino acids, codon models, discrete and continuous morphological features, language, microsatellites and SNPs as well as user-defined discrete and biogeographical data.

Bayesian inference allows the incorporation of many sources of information in the same analysis, such as DNA sequences from extant and extinct species, combined with information from the fossil record. Apart from inferring rooted time trees, which are valuable in and of themselves [10], BEAST also allows addressing many kinds of micro- and macroevolutionary questions, such as determining the age and location of the origin of species and cultures, rates of mutation and migration, and rate of spread of epidemics.

### New BEAST functionality

At the core of BEAST is its MCMC sampling mechanism. This mechanism has been improved for better performance, which is especially useful for analyses with a large number of taxa but little data, such as a geography-only analysis. The calculation time of Felsenstein’s likelihood, i.e., the probability of sequence data given a tree or network and model parameters, which typically takes up the bulk of computing time, has been made more efficient for the case where there is a proportion of invariable sites.

BEAUti has been improved so as to make it easier and more intuitive to set up an analysis. For example, when many tip or clade calibrations are required, these can now be read from a NEXUS file, which tends to be easier to manage than editing calibrations one by one in a GUI. BEAUti now also allows specification of custom tree models, such as multi-monophyletic constraints with multifurcating trees in Newick format as well as switching top-level analyses from MCMC to nested sampling, for example.

While the core of BEAST 2 provides basic functionality for Bayesian phylogenetic analyses, it is mostly a platform for building packages on. Package management has matured to include a command line as well as graphical user interface that can deal with different package repositories. Different versions of packages can be installed at the same time. This is as practical as it is important for reproducibility, because an analysis specification file (the BEAST XML file) generated using an older package version can still be run using that older version without the usual necessity of uninstalling the latest package release. Packages are linked by the GUI to websites, making it easy to find information such as tutorials and user documentation. Packages can also be automatically updated to ensure the latest bug fixes and new features are available.

Finally, BEAST 2 and its tools have been improved and extended to facilitate the implementation of several new packages, which have also been made faster as well as more efficient in their memory usage. The new packages contain most of the new features. In particular, (i) the time trees were extended to generalized phylogenetic structures, (ii) new models for the existing and new structures were developed, (iii) tools for model selections were developed, (iv) and tools for simulating under such models were implemented. We outline these advances in the rest of this paper.

### Beyond time trees: extended phylogenetic structures

BEAST software packages have always dealt exclusively with phylogenetic trees that have an explicit time dimension. The developers of BEAST (and some other Bayesian phylogenetics packages) have championed the notion that time is a fundamental dimension to connect independent sources of evidence about evolution and ancestry; in other words, all evolutionary hypotheses should have the time dimension as an explicit part of their parameterisation. The attraction of doing so is manifold, and has been the primary means by which different quantitative theories from phylo- and population genetics have been melded together into increasingly sophisticated hierarchical phylogenetic models that are now starting to be more regularly employed.

The ancestral structures estimated by BEAST all have a time dimension, but they are not all the classic binary rooted time trees with samples at the tips. Generalizations of a binary rooted time tree structure (Fig. 1a) are essential in certain cases, for example:

**Fig 1.**
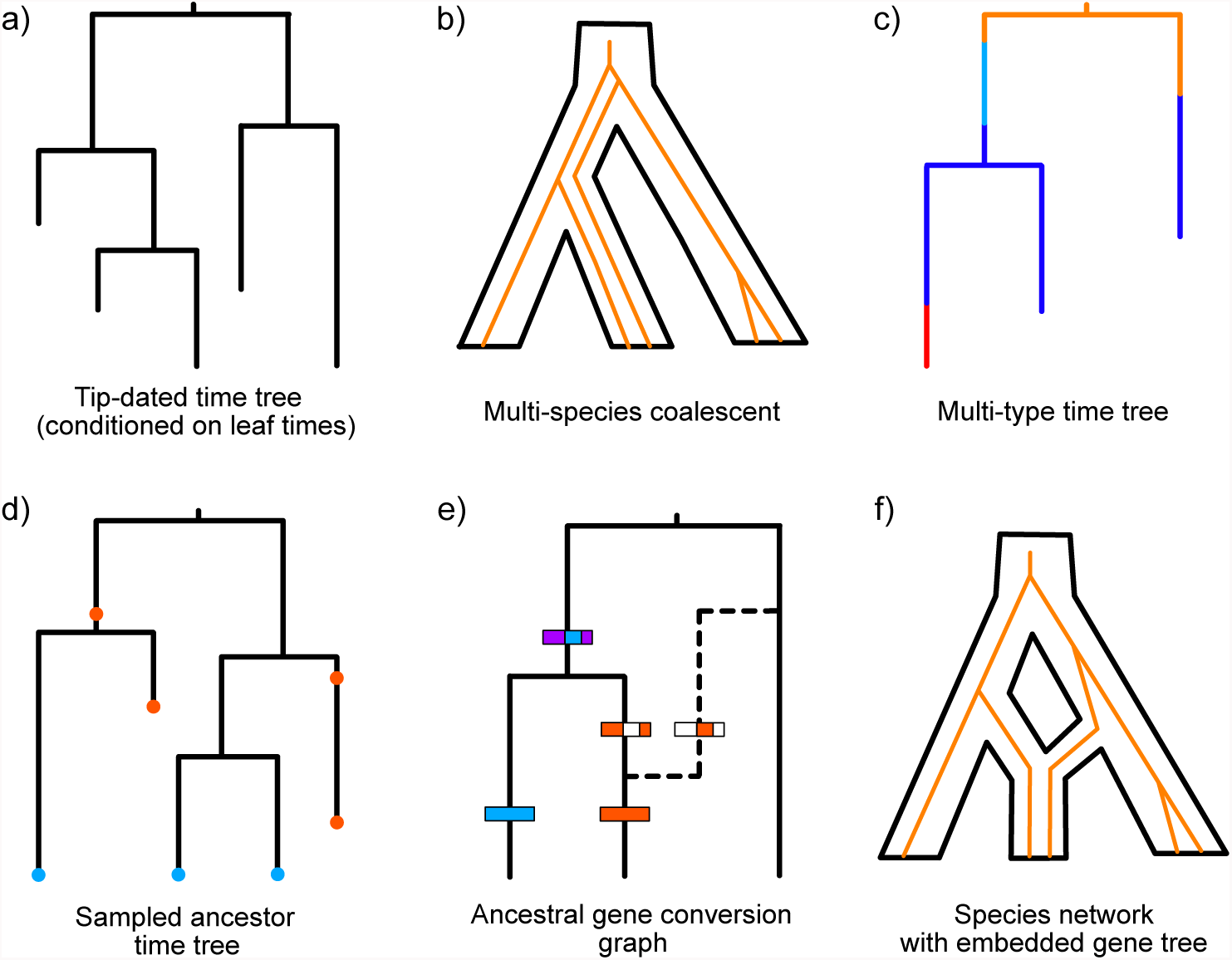
Phylogenetic structures available in BEAST 2. (a) A tip-dated time tree, with leaf times as boundary conditions but not data (generally a coalescent prior is applied in this setting). (b) A species tree with one or more embedded gene trees (c) A multi-type time tree has measured types at the leaves and the type changes that paint the ancestral lineages in the tree are sampled as latent variables by MCMC. (d) A sampled ancestor tree, with two types of sampling events: extinct species (red) and extant species (blue). Extinct species can be leaves or, if they are the direct ancestor of another sample, degree-2 sampled ancestor nodes. (e) An ancestral gene conversion graph is composed of a clonal frame (solid time tree) and an extra edge and gene boundaries for each gene conversion event. (f) A species network with one or more embedded gene trees.

- **population and transmission trees**: branches represent not one lineage, but entire populations (or species) [7, 11], and branching events represent population splits (or speciation or transmission events) [12] (Fig. 1b),
- **sampled ancestors**: fossils may be direct ancestors of other fossils or extant species [13] (Fig. 1d),
- **structured populations**: branches are painted according to which population the individual belongs to [5] (Fig. 1c),
- **clonal frame ancestral recombination graph**: some gene regions have alternative parent edges added to a “clonal frame” phylogeny, resulting in a tree-based network [14] (Fig. 1e),
- **species networks**: hybridization or admixture after isolation events are included in the species history (so that the species history is a directed network) but gene histories (genealogies) are still represented by binary trees [15] (Fig. 1f),
- **polytomies**: one individual gives rise to many lineages at the same time.

Since the first release of BEAST 2, a range of Metropolis-Hastings proposal distributions has been developed to sample these extended phylogenetic data structures using MCMC. Additionally, we need to assume a phylogenetic (or “tree”) prior or model for each such phylogenetic structure. This expansion of the space of possible hypotheses that can be addressed by BEAST 2 continues at pace. In the next section, we will highlight the generative priors for the first four classes of extended phylogenetic structures as well as recent advances on new models for classic binary rooted time trees. In addition, some of us (TGV, TS) are currently working on including time tree polytomies in BEAST 2, as may be relevant to, for example, super-spreading events in infectious disease.

### New models

A Bayesian phylodynamic analysis requires the specification of a model for substitutions, a clock model, and a population dynamic model generating the phylogenetic structure, whether that be a tree, a phylogenetic network or a hierarchical combination of the two. These models induce probability distributions for the proposed states of the MCMC, the MCMC samples from the posterior distribution

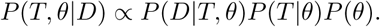

Here *D* is the sequence data and any other sort of data, *T* is the phylogenetic structure as introduced in the previous section, *θ* is the collection of the phylodynamic model parameters, as well as parameters for the substitution, site and branch rate sub-models. The strength of BEAST 2 is that developers can contribute new (sub-)models via packages. Table 1 shows the majority of currently available packages - ordered by their features. An up-to-date list of packages can be seen either from the *Package Manager* embedded in BEAST 2 or using *Package Viewer* (http://compevol.github.io/CBAN/) online.

**Table 1.**
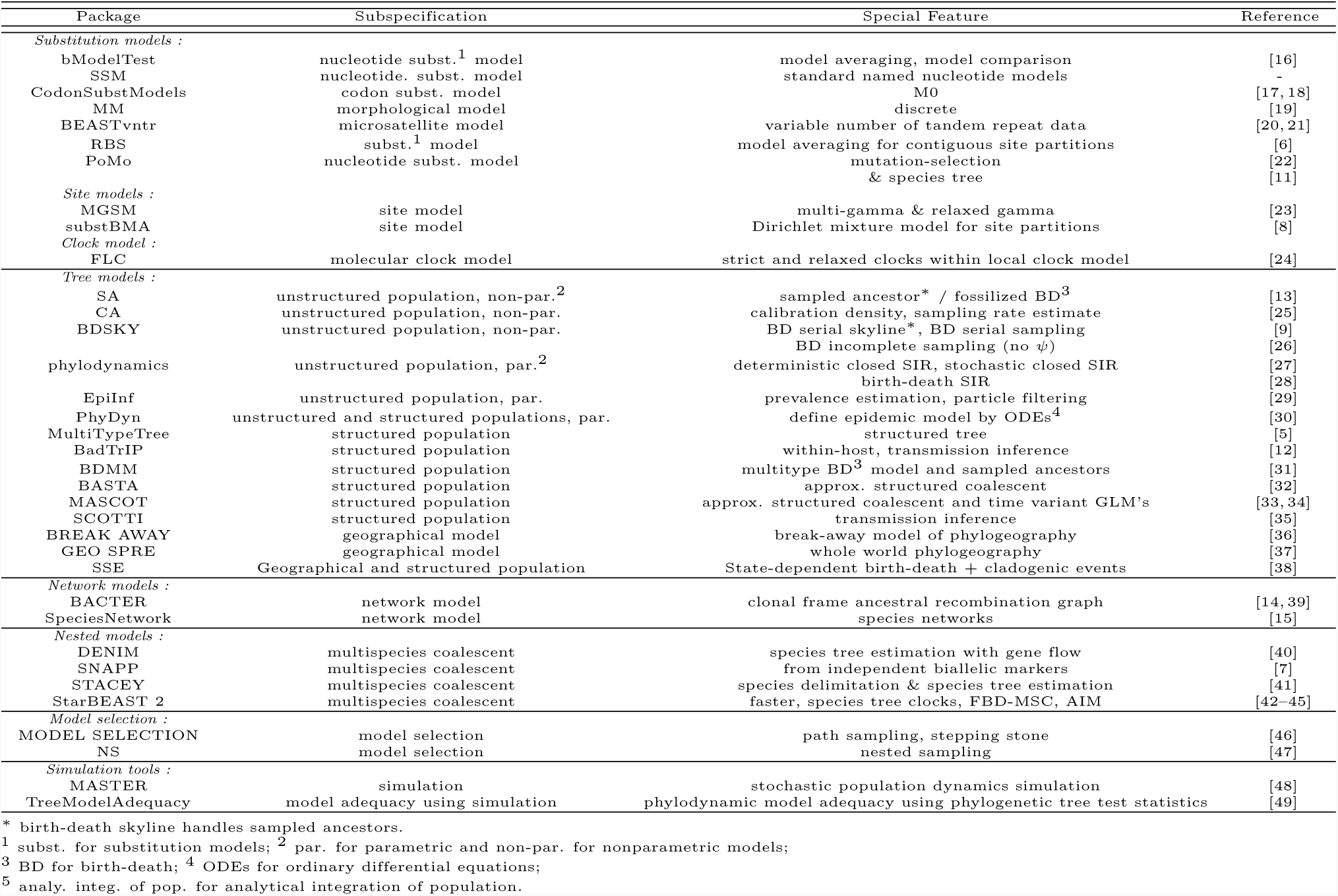
BEAST 2 packages

Below, we highlight some of the key new models in BEAST 2.5, that have been developed since our first description of the BEAST 2 software platform.

### Site models

The site model (encompassing the substitution model and the model rate heterogeneity across sites), together with the clock model, determine the probability *P* (*D|T, θ*) (the phylogenetic likelihood). Model averaging and model comparison of site models are both provided by the new bModelTest package [16]. This package implements reversible-jump MCMC between time-reversible site models for nucleotides, as well as the estimation of the relative support for (i) equal or unequal base frequencies, (ii) uniform or gamma rate heterogeneity across sites, and (iii) zero or non-zero proportion of invariable sites. By providing model averaging of site models within a single MCMC analysis the uncertainty of the site model is integrated out, so that the phylogenetic analysis does not depend on committing to a specific site model. If the site model is not of direct interest, then the posterior distribution on site models can be ignored (knowing it has been model-averaged); otherwise, if the site model is of interest, then bModelTest provides a posterior distribution over site models, so that a credible set of site models can be constructed, and all pairs of site models can be compared for relative support *a posteriori*.

Figure 2 shows the posterior distribution resulting from a bModelTest analysis of substitution models for 906 nucleotides of cytochrome oxidase II and cytochrome b of 36 mammalian species [50] (for details see http://www.doi.org/10.5281/zenodo.1475369). Each circle represents a substitution model indicated by a six digit number corresponding to the six rates of reversible substitution models (see Figure 2 caption for more details).

**Fig 2.** bModelTest analysis for 36 mammalian species [50]. a) Posterior distribution of substitution models. Each circle represents a substitution model indicated by a six digit number corresponding to the six rates of reversible substitution models. In alphabetical order, these are A! C, A! G, A! T, C! G, C! T, and G! T, which can be shared in groups. The six digit numbers indicate these groupings, for example 121121 indicates the HKY model, which has shared rates for transitions and shared rates for transversions. Here, only models are considered that are reversible and do not share transition and transversion rates (with the exception of the Jukes Cantor model). Other substitution model sets are available. Links between substitution models indicate possible jumps during the MCMC chain from simpler (tail of arrow) to more complex (head of arrow) models and back. There is no single preferred substitution model for this data, as the posterior probability is spread over a number of alternative substitution models. Blue circles indicate the eight models contained in the 95% credible set, models with red circles are outside of this set, and models without circles have neglegible support. b) Posterior tree distribution resulting from the bModelTest analysis.

Other substitution and site models added are PoMo [11, 22] (which can account for within-species variation and GC-biased gene conversion), pseudo Dollo [51], codon models [17, 52], standard named nucleotide models (SSM package), standard empirical amino acid models (OBAMA package), morphological models (MM package) [19] and microsatellite models (BEASTvntr package) [21].

### Molecular clock models

The core BEAST 2 package already provides the relaxed [53] and random local [54] clock models to model substitution rate heterogeneity along a phylogeny. The FLC [24] package provides a framework that integrates the flexibility of the relaxed clock model into the local clock model. Specifically, the FLC model allows a local clock to be either strict (i.e. as in the original local model definition) or relaxed. In practice, this means closely related lineages can be modelled with a single constant rate substitution model (i.e. strict clock model) while other lineages with significant rate variation can be described more accurately with a relaxed clock model. As in the original formulation of the local clock model, the user needs to define the location of the local clock *a priori*.

### Population dynamic models for trees

Population dynamic models provide the probability density of the phylogeny given the parameters, *P* (*T|θ*). Population dynamic models giving rise to phylogenies are also called phylodynamic models.

#### Tree models for unstructured populations

There are two common approaches for modelling the phylogenetic tree, or the genealogy, in phylogenetic inference. The first assumes a classic population dynamic model, namely the birth-death model [55,56], to model the growth of a tree. In a population dynamic birth-death model, through time, each individual gives rise to one additional offspring with rate *λ* and dies with rate *μ*. As we only analyse a fraction of individuals arising in this process, it is necessary to model the sampling process for tips of a birth-death tree. For a variety of simple partially-sampled birth-death trees, the distribution of branch lengths has been derived exactly [57].

Alternatively, a mathematical model for trees known as the coalescent [58, 59] can be used to parameterize the tree in terms of the effective size of the background population, and changes in this effective population size through time. One can interpret the effective population size and its changes as birth-death parameters when making some coalescent approximations [30]. Partially-sampled birth-death models do not make the approximations that coalescent models do, but they depend on a model of the sampling process, and simple sampling models may not always be an adequate description of real data sets. It is an ongoing debate and topic of research to investigate the consequences of coalescent approximations and sampling model assumptions.

Coalescent approaches have been embedded within BEAST since its genesis [60, 61]. Thus, we will not further discuss the basic coalescent approach here. In what follows, we will introduce the basic birth-death models which underwent major development in recent years. Then, we discuss the more sophisticated birth-death and coalescent approaches side by side.

In birth-death models, it is assumed that the first individual appears at some time *t*_0_ before the present. Through time, each individual gives rise to one additional offspring with rate *λ* and dies with rate *μ*. An individual is sampled (e.g. the pathogen of an infected individual is sequenced, or ancient DNA for an individual is sequenced; or a fossil is observed) with rate *ψ*. Upon sampling, we assume that the individual representing the sample is removed from the population with probability *r*.In the case of infectious diseases, *r* is the probability of being cured or treated, such that the individual is not infectious any more upon sampling. In the case of species, we typically assume *r* = 0 as the species continues to exist upon sampling of a fossil. At the end of the process, each extant individual is sampled with probability *ρ*. The probability of a tree (Fig. 1d), given parameters *t*_0_*,λ,μ,ψ,r,ρ* has been derived in [57] for *r* =0, and generalized for *r* ϵ[0, 1] in [62]. A value *r<* 1 necessitates using an MCMC algorithm capable of producing trees with sampled ancestors. Such an algorithm is provided in BEAST 2 via the SA (sampled ancestor) package [13].

This basic model has been extended to account for changes of parameters through time within the bdsky package [9]. In bdsky, time is divided up into one or more intervals, inside of which parameters are held constant but between which parameters may be completely different (i.e. the change of parameters occurs in a non-parametric way).

In epidemiological investigations the birth-death model can be reparameterised by setting the rate of becoming noninfectious, *δ* = *μ* +*ψr* (the total rate at which lineages are removed), the effective reproductive number, *R_e_* = *λ/δ*, and the sampling proportion *p* = *ψ/δ* (the proportion of removed lineages that are sampled). Fig. 3 shows the posterior estimates from a bdsky analysis of the 2013–2016 West African Ebola epidemic. Estimates are based on the coding regions of 811 sequences sampled through October 24, 2015, representing more than 2.5% of known cases. There is evidence that hospital-based transmission and unsafe burials contributed infections to the epidemic [64], thus the SA (sampled ancestor) package was used to account for some percentage of patients continuing to transmit the virus after being sampled (by allowing *r* to be less than 1). *R_e_* was allowed to change over 20 time intervals, equally-spaced between the origin of the epidemic (*t*_0_) and the time of the most recent sample, while the sampling proportion was estimated for every month from March 2014 onwards (when an Ebola virus disease outbreak was declared and the first samples collected). The estimated origin time of the epidemic coincides with the onset of symptoms in the suspected index case on December 26, 2013 [63]. Estimates of *R_e_* are consistent with WHO estimates [65], based on surveillance data alone, but with greater uncertainty. For the majority of the period between mid-May and October 2014 *R_e_* is estimated to be above 1, consistent with the observation that September 2014 was the turning point of the epidemic and that case incidence stopped growing in October [65]. After peak incidence was reached during the last week of September 2014, *R_e_* estimates drop below 1 during October and November 2014 and then fluctuate around 1 during 2015 as transmissions persisted in some areas, due to a combination of unwillingness to seek medical care, unsafe burials and imperfect quarantine measures [63]. *R_e_* estimates before May 2014 and after August 2015 have a large amount of uncertainty attached to them, due to the small amount of sequences sampled during these time periods. Trends in sampling proportion estimates follow empirical estimates based on the number of confirmed cases; however, the sampling proportion is overestimated during the period of intense transmission, which suggests the existence of transmission chains not represented in the sequence dataset. In the final two months of the study period the sampling proportion is underestimated, which may indicate ongoing cryptic transmission during this period, but may also be indicative of a model bias resulting from the remaining transmission chains at this time being highly isolated from each other, which is not taken into account by the model.

**Fig 3.**
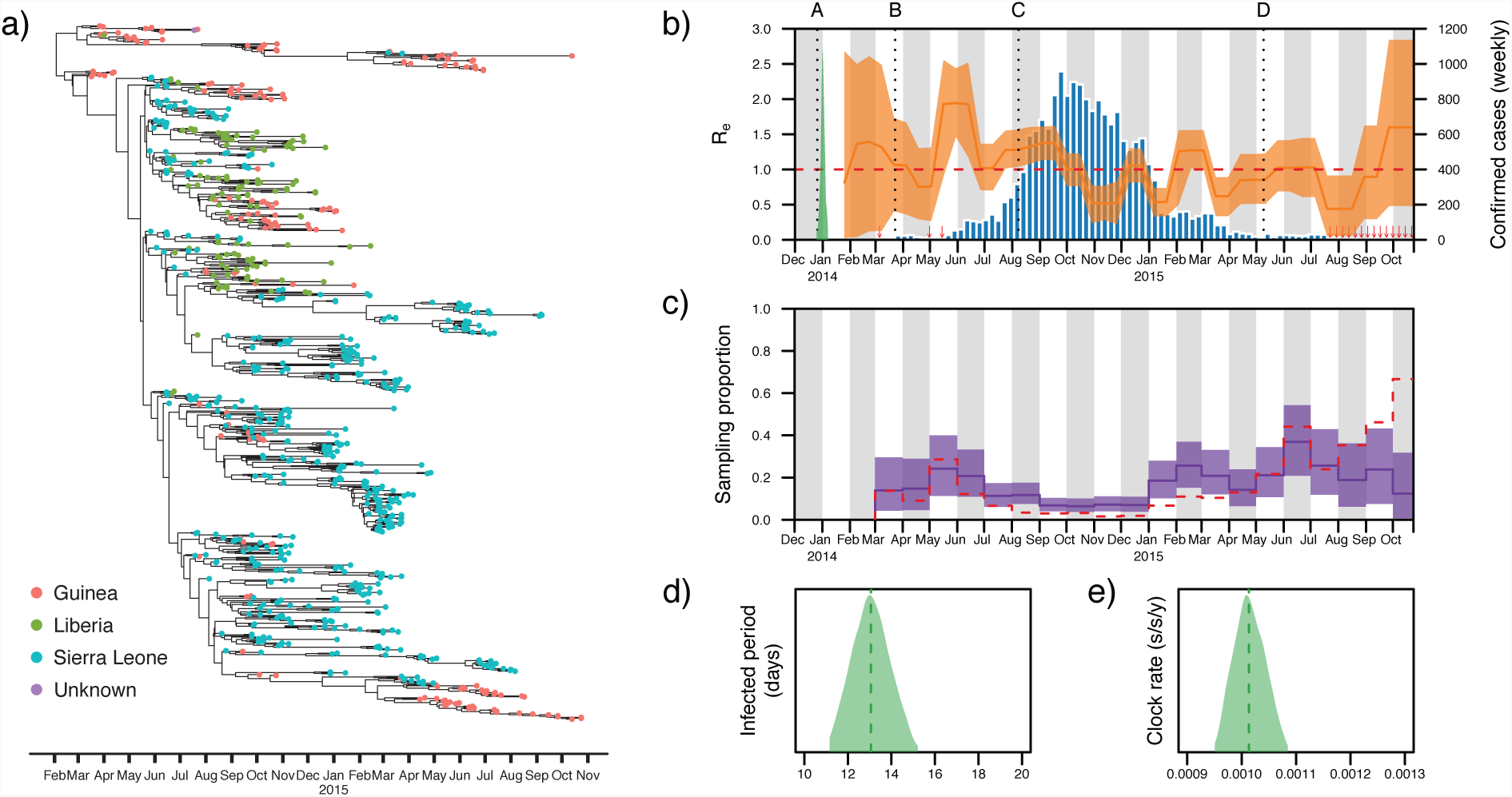
Birth-death skyline (bdsky) analysis of the 2013–2016 West African Ebola virus disease epidemic. (a) The maximum clade credibility tree of the 811 sequences used in the analysis. (b) The median posterior estimate of the estimated effective reproductive number (*R_e_*) over time is shown in orange, with the 95% highest posterior density (HPD) interval in orange shading. The red dotted line indicates the epidemic threshold (*R_e_* =1). If *R_e_*is below this threshold the epidemic has reached a turning point and is no longer spreading. The posterior distribution of the origin time of the epidemic (*t*_0_) is shown in green. The number of laboratory-confirmed cases per week is shown in blue. Red arrows indicate weeks with fewer than 10 confirmed cases. The dotted line at A indicates the onset of symptoms in the suspected index case [63]. The dotted lines at B and C indicate the dates at which the WHO declared an Ebola virus disease outbreak in Guinea and a Public Health Emergency of International Concern (PHEIC), respectively. The dotted line at D indicates the first time any of the three countries with intense transmission (Liberia) was declared Ebola free following 42 days without any new infections being reported (new cases were subsequently detected in Liberia in June 2015). (c) The median posterior estimate of the monthly sampling proportion is shown in purple, with the 95% HPD interval in purple shading. The red dashed line indicates the number of sampled sequences in the dataset, divided by the number of laboratory-confirmed cases, for each month in the analysis. This serves as an empirical estimate of the true sampling proportion. The posterior distributions and medians (dashed lines) of the infected period and the mean clock rate (truncated at the 95% HPD limits) are shown in panels (d) and (e).

Popular models in epidemiology, such as the SIR model [66], or in macroevolution, such as the diversity-dependent model [67], assume that parameters change as a function of the number of susceptible individuals or non-occupied niches, for example. Thus, they are called parametric birth-death models. Such parametric rate changes can be assumed when using the EpiInf package [29]. This latter package additionally samples the trajectory of infectious and susceptible individuals through time and allows for the inclusion of case count data in addition to sequences. In a faster, but approximate way, the phylodynamics package [28] performs inference under the SIR model using genetic sequences.

Parametric birth-death-based population dynamic models are computationally expensive because parameters are a function of the number of co-occurring individuals: typically we do not know this number and thus have to sample it via MCMC. An alternative is to approximate the population dynamics using the coalescent, which essentially means that we assume that our sample is small within a large population, and that we condition on the sampling times instead of them being part of the data, as in the birth-death model. The phylodynamics package provides an approach to estimate the trees and parameters assuming an either deterministically or stochastically changing population size under an SIR-type coalescent framework [27].

The analysis of genetic data and fossils for reconstructing a species phylogeny can be achieved using the birth-death model when setting *r* = 0. This setting is also referred to as the fossilized birth-death (FBD) process [68–70]. These approaches generalize the total-evidence dating method [71, 72] by allowing for sampled ancestor fossils (instead of assuming all fossils are tips in the tree) and modelling of the fossil sampling process. These FBD approaches are an alternative to dating phylogenies by node-calibration approaches. Some constructions of the latter result in complex marginal priors for calibrated nodes [73], and it is not straightforward to specify a prior distribution for each calibration node. Furthermore, node-calibration approaches do not coherently use all comparative data within a joint inference framework, since the decision of which node to calibrate with which fossil is made before phylogenetic inference. This incoherency is overcome by total-evidence approaches where all data is analyzed together and node ages and tree topology are estimated jointly. On the other hand, the FBD models use each fossil age as an observation, and can be very sensitive to a biased fossil or extant species sampling [69, 74]. This is particularly problematic when only the oldest fossils of clades are included in the analysis, as is commonly done in node dating approaches. I such cases, the CA (CladeAge) [25] package allows unbiased age estimation; however, it requires that sampling parameters are known *a priori* of the analysis while the FBD approach estimates these parameters alongside the tree. On the other hand, this requirement of the CladeAge approach means that different sampling parameters can be specified for different clades, whereas all (coexisting) species are assumed to share the same sampling parameters in the FBD model.

### Tree models for structured populations

Methods for studying population structure and reconstructing migration history have seen considerable progress in recent years, and have been particularly bolstered by the modularity and extensibility of BEAST 2. These features represent a remarkable opportunity for end users, who can now use, test and compare different models and approaches without the need to switch platforms and formats. It also encourages method development, as the availability of packages in a single, modular platform aids future development through easy integration of ideas and code.

In analogy with the situation for unstructured populations, the two approaches for structured populations are (i) multi-state birth-death models [9], implemented in the bdmm [31] package, and (ii) structured coalescent approaches, with an exact implementation available within MultiTypeTree [5]. The birth-death and coalescent approaches from above are essentially generalized to allow for more than one population by assuming migration rates between, and variable birth rates across, populations.

The bdmm package allows for changes in dynamics through time by using a skyline, analogous to the unstructured birth-death models. Furthermore, it can quantify its parameters, such as migration rates, without MCMC sampling of the states in ancestral lineages. In other words, for *T* being a phylogenetic tree with its tips being assigned states, bdmm uses equations for *P* (*T|θ*) under the multi-state birth-death model. The bdmm functionality was recently extended for macroevolutionary trees through the SSE package [38]. This package implements a family of (birth-death) models of state-dependent speciation and extinction ranging back to the original BiSSE model [75] where all tips are sampled at one point in time. The “state” a species or population is in can represent the state of one of its traits, but it can also be seen as its geographical distribution. When inputs are geographical ranges, state transition parameters can be interpreted as migration rates.

For the structured coalescent, the MultiTypeTree package samples the ancestral states of all lineages (Fig. 1c), using MCMC, which can become very slow (i.e. MultiTypeTree considers *P* (*T|θ*) with *T* being a phylogeny where all lineages at all times have states assigned). Furthermore, the package needs to assume constant population sizes through time for the different demes. These limitations have been overcome by tracking ancestral states probabilistically using different approximations [30, 76], avoiding the need to sample ancestral states using MCMC. The approximation originally proposed by [30] tracks state probabilities assuming that the state of each lineage evolves completely independently of other lineages in the phylogeny. Thus, an approximate equation for *P* (*T|θ*) under the structured coalescent is employed, where *T* is a phylogenetic tree, with its tips being assigned states. BASTA [32] implements a highly optimized version of the approach of [30] in BEAST 2.5, allowing one to rapidly analyse scenarios with many different sub-populations.

MASCOT [33] implements an improved approximation, derived in [76], that is more closely related to the exact structured coalescent, in that lineage state probabilities reflect the likelihood of each lineage coalescing with other lineages based on their probable location. Simulations using MASCOT revealed no biases in the estimates of parameters and node locations [76]. MASCOT additionally allows estimates of migration rates and effective population sizes across different sub-populations and time to be informed from predictor data (such as clinical, demographic, or behavioural variables) using a generalized linear model (GLM) approach [34].

The PhyDyn package [77] supports a highly flexible mark-up language for defining demographic or epidemiological processes as a system of ordinary differential equations. PhyDyn implements three approximations of the structured coalescent and extended previous work [30] to improve accuracy and reduce computational cost. The package calculates migration and coalescent rates from population trajectories and uses the structured coalescent approximations to calculate the states of lineages through time. A suitable application for this approach is the estimation of parameters from complex infectious disease models with multiple compartments, and it provides a means of taking advantage of categorical metadata which is not related to geography, such as clinical, demographic, or behavioural variables in phylodynamic studies of infectious disease dynamics.

These coalescent frameworks in BEAST 2.5 extend earlier developments on the coalescent. Among the most popular earlier models of this class for studying migration, spread and structure were the structured coalescent-based methods of Migrate-n [78]. Migrate-n targets the same structured coalescent distribution as MultiTypeTree, but differs with respect to the exact implementation. In particular, since not embedded within BEAST, it cannot be coupled with e.g. relaxed clock models.

The very popular discrete trait model and continuous phylogeographic methods from Lemey and colleagues [79, 80] assume that the whole tree was generated under an unstructured model, and that the trait evolved—just like a nucleotide—on that tree. This approach is extremely computationally efficient and allows the study of a large number of samples with many distinct trait values. However, these models make strong assumptions about the distribution of sampled trait values which can bias inference results [32]. This issue can be overcome by the newer but computationally more demanding methods above. The Lemey et al. models are available in BEAST 2 through the beast-classic package (except for the generalized linear model feature introduced in [81]).

Another class of models of population structure deals with the fact that each host in an outbreak contains a separate within-host pathogen population during colonisation. In this context, transmission between hosts is a migration event into a new deme that is consequently colonised. The common aim of such models is to reconstruct the series of transmission events between hosts that led to the establishment of the considered outbreak. BEAST 2.5 offers two different models of such dynamics; SCOTTI [35] models transmission in a structured coalescent setting, and assumes that there is no recombination, that transmission inocula are small, and that each sample consists of an individual haplotype (however, multiple samples from the same host are allowed). BadTrIP [12] instead models transmission with a multispecies coalescent (MSC) paradigm, allowing recombination, large transmission inocula, and within-sample pathogen genetic diversity information from read-based allele counts, while accounting for sequencing error. BadTrIP can efficiently utilize information from genetic variation within samples to reconstruct more detailed transmission histories than SCOTTI, but it is also more computationally demanding [12].

### Multispecies coalescent models

The multispecies coalescent (MSC) model describes the evolution of genes within species [82]. Broadly, it assumes that the sampled alleles for a given gene have evolved according to a common coalescent process within each species, typically thought of as occurring backwards in time. For each branch in the species tree, this process begins at the tipward end of the branch, and apart from the root is truncated by the speciation event at the rootward end. Thus the MSC models trees within trees, and the probability density *P* (*T|θ*) becomes more complex, as described below.

An emergent property of the MSC known as incomplete lineage sorting (ILS) occurs when two or more lineages do not coalesce in their immediate ancestral population (Figure 4), which can lead to gene trees with discordant topologies among themselves and with the species tree. The probability of ILS increases as branch lengths are shortened in time, and/or when the effective population size *N_e_* is increased. Species trees with four or more ingroup species can have a region of their parameter space (the “anomaly zone” [83]) where most gene trees have a topology different to the one of the species tree.

**Fig 4.**
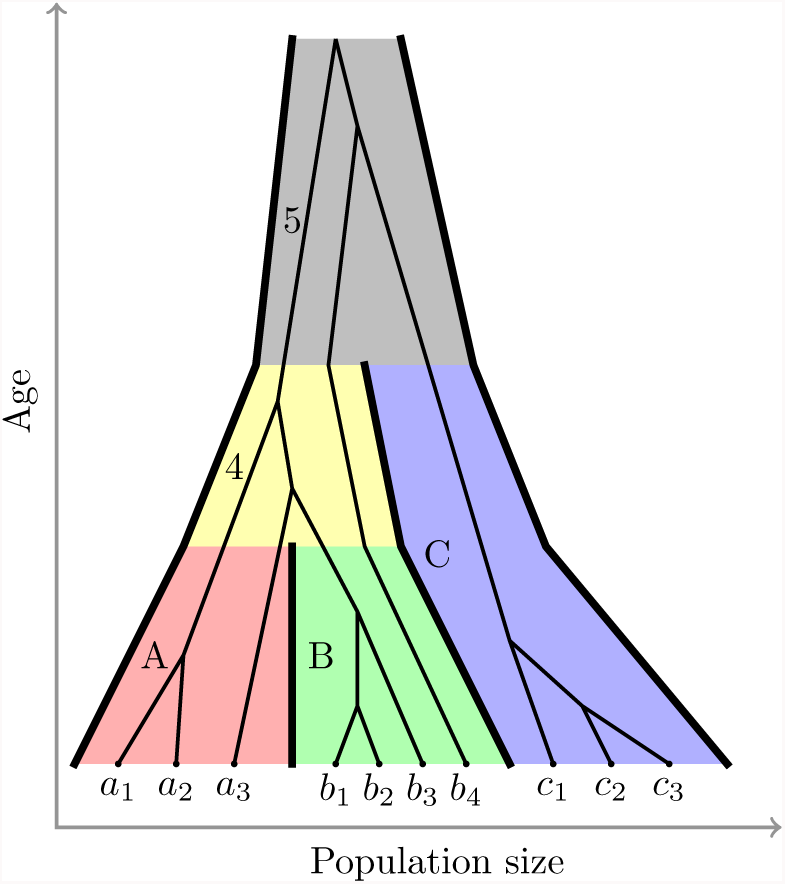
The multispecies coalescent (MSC) model with three species and a single gene tree. A separate coalescent process applies to each of the five branches in the tree; the branches for the extant species A (red), B (green) and C (blue), the ancestral branch of A and B (yellow), and the root branch (grey). Several individuals have been sampled per species. In this example the ancestral lineage of individual *b*_4_ does not coalesce in species B or ancestral species 4. In ancestral species 5, it coalesces with the ancestral lineage of species C. This leads to incomplete lineage sorting and enables gene tree discordance – in this example *b*_4_ is a sister taxon to individuals from species C, rather than to individuals from its own species, or sister species A. If *b*_4_ was the representative individual for its species, then this gene would exhibit gene tree discordance. Other individuals which show concordance at this locus are expected to show discordance at other unlinked loci when populations are large or speciation times are recent.

Discordance between gene trees and species tree in their topologies and times can lead to incorrect species tree estimates from concatenated gene sequences – this has been shown to occur with both maximum likelihood and Bayesian methods like those implemented in BEAST. More specifically, in the anomaly zone, gene tree topological discordance can result in incorrect estimates of the species tree topology [84, 85], and systematic bias in branch length estimates [86]. Even in the case of just two species where gene tree discordance is impossible, speciation times estimated using concatenation will be wrong because the expected time to coalescence is 2*N_e_* generations older than the speciation time [87]. The concatenation estimates of speciation times are therefore expected to be 2*N_e_* generations older than the truth.

Unlike concatenation, multilocus MSC methods can accurately and jointly estimate the topology and times of the species tree and gene trees directly from multiple sequence alignments (MSAs). The first BEAST multilocus MSC implementation was *BEAST, which was introduced in BEAST 1.5.1 [88]. Let *P*(*T,G,θ*|*D*) be the joint posterior probability density for a species tree (*T*), a set of gene trees (*G* = *g*_1_*, g*_2_*,…, g_L_*) and additional evolutionary parameters (*θ*), given a corresponding set of multiple sequence alignments *D* = {*d*_1_*, d*_2_*,…, d_L_*}. Thus, we now enrich our posterior probability from above, *P* (*T, θ*|*D*) by additionally sampling gene trees *G*, using *P* (*T, G,θ*|*D*). In the MCMC, we calculate the product of phylogenetic likelihoods *P* (*D_i_*|*g_i_, θ*), the coalescent probability density *P* (*g_i_*|*T, θ*) for each gene tree *g_i_*, and the prior probability of the species tree given macroevolutionary parameters *P* (*T* |*θ*):

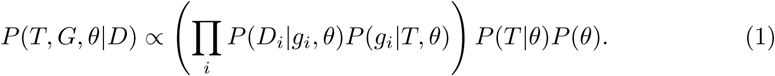

StarBEAST 2 [43] built on *BEAST [88] introduced species tree relaxed molecular clocks, where a separate substitution rate is estimated for each branch of the species tree. The substitution rates across each gene tree, used to calculate gene tree likelihoods, are then derived from the per-species rates and the per-gene rates [43]. This clock model enables accurate inference of substitution rate variation across the species tree from multiple loci.

Recently, some of us have developed an integrative model of molecular and morphological evolution which combines the FBD and MSC models to infer species trees from neontological and paleontological data, called the FBD-MSC for short. In this model, morphological data evolve along the species tree like the FBD model, but the MSC is used to model molecular evolution. The FBD-MSC was implemented in StarBEAST 2 v14. Using simulation, it was shown that differences in estimated ages between concatenation and the FBD-MSC are likely due to systematic biases introduced by concatenation [44].

Although the MSC deals successfully with a ubiquitous source of discordance, it has limitations. It relies on an assumption that there is no recombination within loci and free recombination between loci. The MSC also ignores the possibility of hybridization. Furthermore, in the MSC, speciation is assumed to be immediate, with an instant where (going back in time) coalescence suddenly becomes possible. In practice, speciation is usually expected to be gradual, and sometimes gene exchange occurs between non-sister species. Newly developed approaches relaxing such strict tree constraints are described in the next section on explicit models of reticulate evolution.

Another assumption of the MSC is that individuals can reliably be assigned to species or populations, whereas in practice, this is often not the case, especially with shallow phylogenies. DISSECT [89], extending the MSC, was first developed for BEAST 1.8.1, and it makes no assumption about how individuals are grouped into species, by inferring species assignment and delimitation simultaneously with the joint inference of the species and gene trees. It does so through an approximation to the Dirac delta function, where the birth-death prior includes an additional probability ‘spike’ of very short duration,*E*, just before the present. This model is called the birth-death-collapse model. When the most recent common ancestor (MRCA) of multiple individuals is present inside the spike, those individuals are often interpreted as belonging to a single species [90, 91].

Improving the computational performance of MSC methods is an ongoing challenge. Increasing the number of individual specimens in an analysis will degrade computational performance. Most seriously, the relationship between the number of loci used with

*BEAST and the time taken to collect enough independent samples from the posterior distribution follows a power law distribution. The result is that whenever the number of loci used in a study is doubled, the time taken to run *BEAST increases seven-fold [42].

Both StarBEAST 2 and STACEY [41] (the successor of DISSECT) offer improved MCMC mixing over their predecessors. STACEY introduced a number of new classes of MCMC operators that simultaneously modify the species and gene trees in a coordinated fashion. On a data set where *BEAST was not able to converge when used with any more than 50 loci, STACEY was successfully run with 500 loci [41].

Likewise StarBEAST 2 has implemented coordinated operators belonging to one of the classes introduced by Jones [41]. Both StarBEAST 2 and STACEY also implement analytical integration of population sizes, which reduces the number of parameters which must be estimated using MCMC. The combination of new operators, analytical integration and additional optimizations to data structures enables StarBEAST 2 to be run with double the number of loci in roughly the same time as *BEAST.

Other approaches have addressed the computational burden associated with the MSC by taking a different modeling path. In particular, it is possible to greatly reduce the number of parameters associated with the gene trees in the MSC by integrating over all possible gene trees at each locus and at each MCMC step. This way, the parameter space does not increase as new loci are added to the analysis, and computational demand increases typically only linearly with the number of loci. In order to simplify gene tree integration, these models consider individual sites as loci, treating each SNP, or base, as unlinked from the others. While this modeling assumption can represent a coarse approximation, it on the other hand has the advantage of allowing recombination within genes, that otherwise can bias gene tree (and therefore species tree) inference.

One of the first gene tree-integrating approaches was SNAPP [7], which infers species trees directly from a matrix of biallelic markers (without linkage between markers), and is available as a package for BEAST 2. SNAPP integrates over all possible gene trees for each marker at each MCMC step, enabling much wider data matrices of thousands of markers to be used. The posterior probability density becomes:

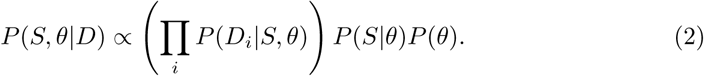

Another similar approach is PoMo [11]. PoMo models each species in the species tree as a small population (in particular, a Moran model [92]), affected by new mutations (introducing new low-frequency alleles in a population) and genetic drift (changing allele frequencies within populations). Differently from SNAPP, PoMo uses nucleotide data, allowing more than two alleles at each SNP, but still allowing at most 2 alleles at one time at any species/population. For each species and locus, PoMo reads 4 numbers, corresponding to the allele counts of the 4 nucleotides at the considered species and locus. PoMo is generally faster than SNAPP or MSC methods [11], and in its BEAST 2 implementation it can account for sequencing errors, as for allele counts derived from reads mapped to a reference genome.

### Reticulate evolution

Describing evolutionary history using tree structures is generally a simplification. Genomes are subject to recombination, organisms are subject to horizontal gene transfer and species undergo hybridization followed by introgression. With a small number of exceptions (e.g. [93], [94]), computational phylogenetics has so far addressed these processes only partially, by restricting gene tree reconstructions to relatively short alignments that are assumed to be free from intra-locus recombination, or by excluding taxa from phylogenetic analyses that were found to be involved in gene flow by other approaches [95].

However, while these approaches to some extent avoid bias resulting from recombination, they at the same time ignore it as a potentially very useful source of information that is increasingly provided by whole-genome sequencing. For example, it has been shown that making use of this large-scale genomic structure can lead directly to powerful insights into ancestral population dynamics [96, 97]. Similarly, with the increasing sophistication of species history reconstruction methods brought about through the availability of MSC methods, the omission of important processes such as hybridization and horizontal gene transfer from these models is becoming obvious. In response to this demand, BEAST 2 package authors have contributed and/or implemented a number of algorithms which perform phylogenetic/phylodynamic inference under models which directly account for non-tree-like evolution.

### Gene conversion

The package Bacter [14] provides a complete, carefully validated, reimplementation of the ClonalOrigin model [39] which approximately describes networks produced by homologous gene conversion in bacteria. This is done by approximating the recombination graph using a tree-based network [98], in which the underlying tree is the “clonal frame” produced by the bacterial reproduction process and the additional edges represent homologous gene conversion events. In contrast to the original implementation, BACTER allows for joint estimation of both the clonal frame and the reticulations contributed by conversion events. Additionally BACTER provides a heuristic algorithm for summarizing the posterior distribution over these trees in a fashion similar to the MCC tree approach used by BEAST for binary trees.

### Hybridization and horizontal gene transfer

For multispecies phylogenetic analyses, a model called the Multispecies Network Coalescent (MSNC) has been developed [99, 100]. This model generalizes the MSC by replacing the species tree (which supports only speciation nodes) with a species network (supporting speciation and reticulation nodes). Reticulation nodes and edges in the network can represent multiple biological processes including hybrid species, introgression or secondary contact. Gene trees, embedded within the species network, are still used to model the evolution of individual loci. This means the MSC’s assumption of no intra-locus recombination still applies.

SpeciesNetwork, a fully Bayesian implementation of the MSNC where the species network and gene trees are estimated directly from MSAs, has been developed and is available as a package in BEAST 2.5 [15]. Unlike for the MSC, there may be more than one possible embedding of a gene tree of given topology and times within a species network of given topology and times. The probability density of a possible embedding thus depends on the inheritance probability *γ* at each reticulation node.

In SpeciesNetwork, the gene tree embeddings, Ψ, and inheritance probabilities, *γ*Γ, are jointly estimated alongside the species network, gene trees and other parameters. The posterior probability density for the model is similar to *BEAST and StarBEAST 2, but *T* represents a species network rather than a tree, and the additional jointly estimated parameters are included:

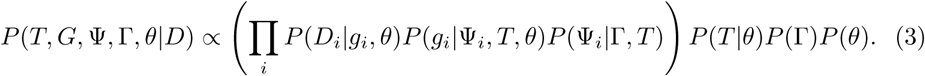

### Isolation with migration

Sitting between the MSC and the MSNC are models where there is a species tree (not network) but the exchange of genes is allowed between the branches of the species tree. This exchange of genes is typically termed gene flow. Gene flow may occur between sister species, known as isolation-with-migration (IM) [101] and between non-sister species (paraphyly) [102]. It has been shown that ignoring gene flow can result in poor estimates of species tree topologies and node times [102].

One solution in the BEAST2 framework is the DENIM package [40], which is able to infer species trees more accurately than MSC-based models such as STACEY when a small amount of gene flow is present. It uses an approximation which breaks down if there is too much gene flow. DENIM is also able to identify which loci are subject to gene flow.

Another solution is AIM [45], which is part of StarBEAST 2 since version v15. AIM implements an IM model that allows the estimation of species trees, rates of gene flow and effective population sizes from genetic sequence data of independently evolving loci. Inferring the species tree topology alongside the other parameters of interest is possible due to the ability to integrate over migration histories [76]. For every set of effective population sizes of extinct and extant species and rates of gene flow between these species, AIM can calculate the probability of a gene tree given a species tree without inferring the migration events. This allows changing the species tree topology and node order while still computing the probability of gene trees under these new settings. MCMC can thus be used to explore the different combinations of species trees, rates of gene flow, effective population sizes and gene trees jointly.

Figure 5 shows the species tree and migration events inferred with AIM from a set of 100 nuclear gene sequence alignments for five species of Princess cichlid fishes(*Neolamprologus savoryi*-complex [103]) from the East African Lake Tanganyika and the outgroup species *Metriaclima zebra* from Lake Malawi. Princess cichlids are well known to hybridize in captivity when placed in the same aquarium [103], and hybridization in their natural habitat has been supported by observed discordance of mitochondrial and nuclear among-species relationships [104]. Whole-genome sequence data for the six species have been generated by [105] and [106] and were used by [106] to generate 426 time-calibrated phylogenies from individual regions of the genomes; a comparison of these phylogenies then supported three past hybridization events in Princess cichlids: between *Neolamprologus brichardi* and *N*. *pulcher*, between *N*. *marunguensis* and the common ancestor of *N*. *pulcher* and *N*. *olivaceous*, and between *N*. *marunguensis* and *N*. *gracilis* [106]. For the analysis shown in Figure 5, we reused the genome data of [105] and [106] to generate alignments for 100 one-to-one orthologous genes following [107], and estimated the species tree jointly with the support for gene flow under the AIM model. We fixed the height of the species tree to be 9.2 Mya [95] and inferred the clock rate and transition/transversion ratio for each locus jointly with all other parameters. The backwards in time rate of gene flow between any two species (except the outgroup) was assumed to be inversely proportional to the time these two species co-existed. For each possible direction of gene flow, we inferred the support for this rate being non-zero [79] and the rate scaler itself. The rate scaler was assumed to be exponentially distributed around 0.05. While not exactly equal, this corresponds in scale to about 5% of lineages to have originated from a different species.

**Fig 5.**
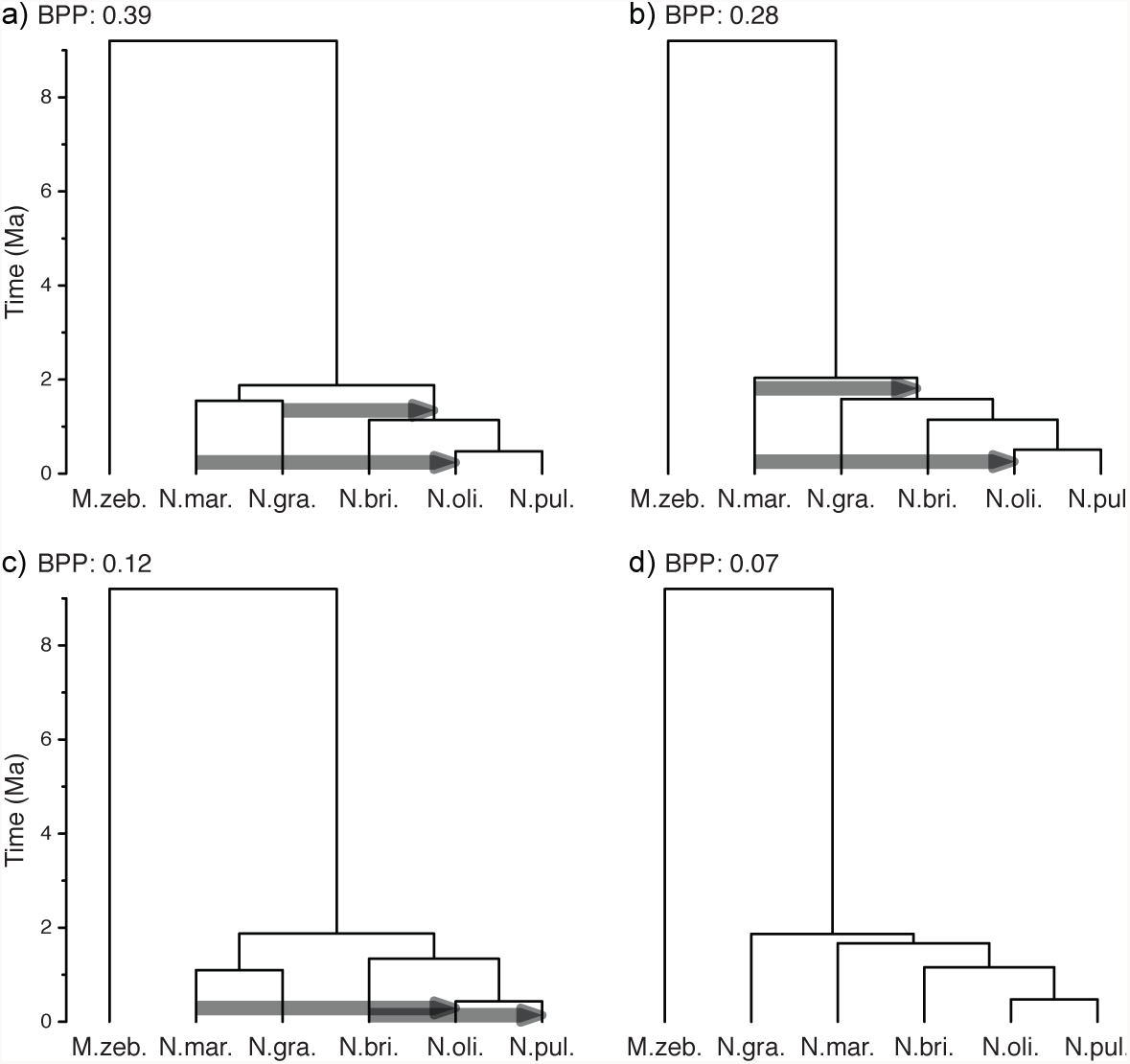
AIM analysis of 100 nuclear gene alignments for the five Princess cichlid species *Neolamprologus marunguensis*, *N*. *gracilis*, *N*. *brichardi*, *N*. *olivaceous*, and *N*. *pulcher*, as well as the outgroup *Metriaclima zebra*. a) to d) show the best-supported tree topologies. Arrows show directions of gene flow that are supported with a Bayes Factor of more than 10. Trees a) and c) only differ in the timing of the speciation events; however, AIM differentiates between differently ranked topologies, since these have to be characterized by using different parameters.

### Model selection and model adequacy

The model selection package has been extended with a number of existing methods, and now contains path sampling, stepping-stone, Akaike information criterion for MCMC (a.k.a. AICM), conditional predictive ordinates [108] and generalized stepping-stone [109].

The NS package implements nested sampling [47] for phylogenetics, which can also be used for model selection. Nested sampling is a general purpose Bayesian method [110] for estimating the marginal likelihood, which conveniently also provides an estimate of the uncertainty of the marginal likelihood estimate. Such uncertainty estimates are not easily available for other methods. Furthermore, nested sampling can be used to provide a posterior sample, and, for some cases where standard MCMC can get stuck in a mode of a multi-modal posterior, nested sampling can produce consistent posterior samples [47]. The marginal likelihood estimates produced by nested sampling can be used to compare models, so provide a basis for model selection.

While model selection compares different models, in model adequacy studies, we assess if a model is a good fit by itself. The key idea of model adequacy assessments is to perform direct simulation of data from generative models (i.e. any of the models discussed above). More precisely, simulations are used to assess the absolute model fit in a posterior predictive framework. First, data is simulated using parameter values sampled from the posterior distribution. Such simulations are known as posterior predictive simulations [111–113]. A test statistic is calculated for the empirical data and for the simulated data. The model is considered to adequately describe the data if the test statistics for the empirical data fall within the range of those from the posterior predictive simulations, for example using a posterior predictive p-value (analogous to the frequentist *p*-value). For example, a phylodynamic model can be used to estimate the reproductive number, the origin of the outbreak, and epidemic trajectories (e.g. [27–29]). The package TreeModelAdequacy (TMA; [49]) can sample the posterior distribution of these parameters to generate trees using MASTER [4] and it calculates a number of test statistics. In Figure 6 we assess the adequacy of stochastic and deterministic phylodynamic models by comparing the root-height of trees generated using posterior predictive simulations for a data set of the 2009 H1N1 influenza pandemic.

**Fig 6.**
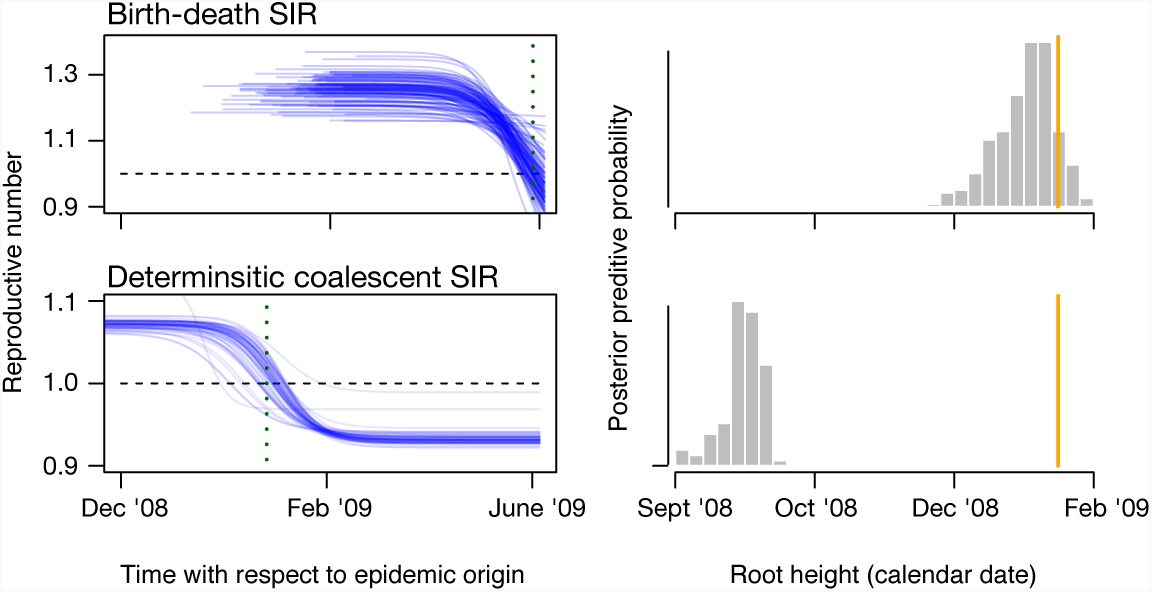
The right column shows the trajectories of the reproductive number over time for a set of 100 publicly available genomes from the 2009 H1N1 influenza pandemic in North America using stochastic (birth-death SIR; [28]) and deterministic (deterministic coalescent SIR [27]) models. Each blue line is a trajectory sampled from the posterior distribution. The models make different inferences of when the reproductive number falls below 1 (vertical dotted line; the horizontal dashed line is for R=1), indicating that the pandemic is past its infectious peak. The right column shows the posterior predictive distributions of the root height for both models (grey histograms) and the value for the empirical data(orange vertical lines). Trees simulated from the stochastic model produce trees that are more consistent with the empirical tree than those from the deterministic model, suggesting that stochasticity may play an important role in the early stages of the pandemic (samples were collected up to June 2009).

## New simulation tools

Many of the models that are implemented in BEAST are generative models that present simplistic, yet mathematically precise, biological hypotheses about the way in which genetic sequences and phylogenetic trees are produced. The focus of BEAST is predominantly learning about biologically meaningful processes via inference of model parameters or model selection. However, models can differ greatly in their assumptions about these processes and the data they generate. Obviously, one must have a clear picture of what generative models imply about data, and if some predicted data features (under a model) are never seen in nature, appropriateness of the model must be questioned. In the previous section, we discussed how to assess model adequacy using simulations.

Furthermore, direct simulation also forms the basis for many inference algorithm validation strategies. Often the best test for correctness of implementation involves judging whether the parameters inferred from data simulated under the model match those used during the simulation. This kind of test can be done qualitatively, or may form the basis for a quantitative validation study by organizing a well-calibrated analysis in which parameters for the data simulation stage are drawn from the same probability distributions used as priors in the inference stage.

BEAST 2.5 provides a number of tools for simulating genetic sequence data and phylogenetic trees. Sequence data simulation is provided as a core feature, and is possible for any of the substitution and clock models supported by BEAST itself or as third-party packages. Phylogenetic tree simulation under specific phylodynamic models (e.g. unstructured/structure coalescent, FBD models, etc.) is provided by the packages that implement those models. General simulation of trees and networks under arbitrary birth-death and coalescent models is provided by MASTER [4], which allows models to be specified using a readable chemical reaction notation and for a wide variety of sampling schemes to be simulated.

BEAST methods have been applied extensively in cultural evolution (e.g., [36, 114, 115]) using the observation that linguistic data can be represented by binary sequence data, and these can be treated similarly to genetic sequence data. The LanguageSequenceGen package [48] can be used to simulate language data under common linguistic models of evolution, with languages specific features like borrowing and burst of evolution shared among different words.

## Availability and Future Directions

BEAST is available under the LGPL licence from https://github.com/CompEvol/BEAST2 and is based on Java, so runs on any platform that supports Java. More information, including downloads, tutorials, news updates, frequently asked questions, etc. can be found on http://BEAST2.org/. Additionally, tutorials for many of the described packages can be found as part of the http://taming-the-beast.org/ platform [116]. At Google groups, there is a forum (https://groups.google.com/forum/#!forum/beast-users) for users to discuss questions.

BEAST 1 is still being developed with a focus on epidemiology of infectious disease, and given its common pedigree it is not surprising that there is considerable overlap in functionality of BEAST 1 and 2. With this in mind, the project X-BEAST (pronounce cross-beast) (https://github.com/rbouckaert/xbeast) is being developed which aims at making two versions of BEAST interoperable, so models from both versions can be used in the same analysis. This non-trivial software engineering problem is something we hope will yield fruit in the near future.

## Discussion and Conclusion

Since the first release of BEAST 2 there has been a large expansion of core features, an increase in the number of developers, and a large increase in the number of models and the number of packages available. There has also been the publication of a book [2] and the introduction of a regular series of week-long in-depth Taming the BEAST workshops [116]. The BEAST 2 community has rapidly grown over the past 5 years and the software has grown (with respect to other similar software packages) in a number of distinct directions: (i) hierarchical multi-species coalescent models for species tree estimation, (ii) fossilized birth-death models for macroevolution and total-evidence analyses and (iii) multi-state birth-death and structured coalescent epidemiological models for understanding rapidly evolving infectious diseases, (iv) new model averaging and model comparison methods including nested sampling. BEAST 2 now occupies a unique niche in the landscape of Bayesian phylogenetic inference software, but still shares a very similar modeling philosophy with both BEAST 1.10 [117] and RevBayes [118]. There are pros and cons to having many different platforms that both compete and complement each other. On the positive side of the ledger, multiple platforms provide the opportunity to validate complex new models by comparing independent implementations. On the negative side, a lack of interoperability means that combining models from two different platforms is currently not possible. So one aim for the future may be to work harder on interoperability between these different platforms. To do so will require a common language for model specification. This is currently the biggest hurdle and an obvious target for future work.

## Supporting information

The XML file and log files used for the bModelTest analyses shown in Fig. 2 are available from http://www.doi.org/10.5281/zenodo.1475369.

The XML file, log file, MCC tree and post-processing scripts for the bdsky analyses shown in Fig. 3 are available from http://www.doi.org/10.5281/zenodo.1476124.

The alignments, XML files, log files and post processing scripts for the AIM analysis shown in Fig. 5 can be found at https://github.com/nicfel/Neolamprologus.

The XML files and a script to generate the TreeModelAdequacy analyses shown in Fig. 6 are available from http://doi.org/10.5281/zenodo.1473852

## Acknowledgements

We would like to acknowledge all additional contributors to BEAST 2 and the Taming the BEAST workshops including Veronika Bošková, David Bryant, Arjun Dhawan, Tracy Heath, Simon Ho, Stéphane Hué, Carsten Magnus, Patricio Maturana, Vladimir Minin, Venelin Mitov, Julija Pečerska, Oliver Pybus, Jérémie Sciré, Christiaan Swanepoel, Erik Volz, Rachel Warnock, David Welch, Jing Yang, Rong Zhang. AJD would like to acknowledge support from a Royal Society of New Zealand Marsden award (#UOA1611; 16-UOA-277). LdP would like to acknowledge support from the European Research Council under the Seventh Framework Programme of the European Commission (PATHPHYLODYN: grant agreement number 614725). Igor Siveroni would like to acknowledge support from the NIH MIDAS U01 GM110749 grant. NFM and TS are funded in part by the Swiss National Science foundation (SNF; grant number CR32I3 166258). TS, JBS, LdP, TGV, and CZ were supported in part by the European Research Council under the Seventh Framework Programme of the European Commission (PhyPD: grant agreement number 335529). DK would like to acknowledge support from the Max Planck Society. MM acknowledges support from the Swiss National Science Foundation (SNP; grant number PBBSP3-138680).

